# p53 engagement is a hallmark of an unfolded protein response in the nucleus of mammalian cells

**DOI:** 10.1101/2024.11.08.622663

**Authors:** Joseph H Park, Thomas J Wandless

**Affiliations:** Department of Chemical and Systems Biology, Stanford University, Stanford, CA, USA

## Abstract

Exposure to exogenous and endogenous stress is associated with the intracellular accumulation of aberrant unfolded and misfolded proteins. In eukaryotic cells, protein homeostasis within membrane-bound organelles is regulated by specialized signaling pathways, with the unfolded protein response in the endoplasmic reticulum serving as a foundational example. Yet, it is unclear if a similar surveillance mechanism exists in the nucleus. Here we leveraged engineered proteins called destabilizing domains to acutely expose mammalian cells to nuclear- or cytosolic- localized unfolded protein. We show that the appearance of unfolded protein in either compartment engages a common transcriptional response associated with the transcription factors Nrf1 and Nrf2. Uniquely, only in the nucleus does unfolded protein activate a robust p53-driven transcriptional response and a transient p53-independent cell cycle delay. These studies highlight the distinct effects of localized protein folding stress and the unique protein quality control environment of the nucleus.

## Introduction

The accumulation of misfolded proteins poses an intracellular threat to cell viability as it disrupts protein homeostasis (proteostasis) and normal biological function (Chen et al., 2011). The accumulation of misfolded proteins and their subsequent transformation into insoluble aggregates is both a causative factor and symptomatic hallmark of many neurodegenerative diseases, such as Alzheimer’s disease, Huntington’s disease, and amyotrophic lateral sclerosis (Hipp et al., 2019). Since proteins can misfold at all stages of their life cycle, all living cells maintain evolutionarily conserved protein quality control (PQC) pathways to ensure chaperone-mediated folding and refolding, sequestration to protective inclusion bodies, and proteolytic degradation of terminally misfolded proteins (Wolff et al., 2014). The ubiquitin-proteasome system (UPS) is a highly conserved pathway in eukaryotes and is the primary proteolytic mechanism that targets both misfolded proteins for degradation as well as mature proteins for metabolic turnover (Ciechanover and Brundin, 2003).

Proteostasis is disrupted when cells encounter stress or transition into pathogenic states. To address these challenges, cells have evolved stress response pathways that engage various proteins that function as sensors and effectors to mount coordinated transcriptional and translational remodeling (Santiago et al., 2020). For example, in response to elevated temperatures that cause protein misfolding and damage to cellular components, the heat shock response (HSR) upregulates molecular chaperone proteins (Richter et al., 2010). Additional PQC pathways exist and have been characterized for several specific types of stress, such as the oxidative stress response (Santiago et al., 2020), integrated stress response (Costa-Mattioli and Walter, 2020), unfolded protein response in the secretory pathway (erUPR) (Hetz et al., 2020), and mitochondrial unfolded protein response (mtUPR) (Shpilka and Haynes, 2018).

One hallmark of eukaryotic cells is subcellular compartmentalization facilitated by membrane-bound organelles (Cohen et al., 2018). These membranes serve as physical barriers that spatially and temporally sequester specialized cellular processes. In some cases, these barriers protect from intrusion by unrelated biochemical processes in addition to intracellular threats (Bar-Peled and Kory, 2022). The nucleus, for example, houses the genome and the nuclear envelope restricts the transport of genetic material, whether it is leakage of genomic DNA to the cytosol or invasion by viral genetic material (Maciejowski and Hatch, 2020; Yang et al., 2023). However, these membrane barriers also necessitate specialized pathways to maintain compartment-specific proteostasis and enable inter-organellar communication to coordinate transcription and translation (Wolff et al., 2014). The erUPR and mitoUPR are two such specialized PQC pathways that respond to acute and chronic stress. Like the ER and mitochondria, the nucleus is a membrane-bound organelle. Although recent work has begun to uncover the distinctive proteostatic challenges and functions of the nucleus (Buggiani et al., 2024), it remains unclear whether a unique nuclear unfolded protein response to acute proteotoxic stress exists.

Our lab previously investigated whether mammalian cells are sensitive to the appearance of a single species of unfolded protein expressed throughout the cell (Miyazaki et al., 2015). When exposed to a single species of unfolded protein, mouse NIH3T3 cells mount a robust, cross-protective transcriptional response. Interestingly, this response engages the p53 pathway. In contrast, these same cells do not engage p53 when transiently exposed to heat shock to induce the HSR or treated with tunicamycin to induce the erUPR. The engagement of p53 by the appearance of unfolded protein, but not by these other protein folding stresses, suggested an unexplored role for p53 in PQC surveillance and regulation. The tumor suppressor p53 is an unstable protein with a half-life less than 20 min and is sensitive to disruptions of the UPS (Pant and Lozano, 2014). The p53 protein stabilizes in response to both specific perturbation (e.g., small molecule inhibition of E3 ligases) and global disruption of the UPS (e.g., proteasome inhibition) (Pant and Lozano, 2014). When stabilized, p53 protein transactivates target genes that induce cell cycle arrest, senescence, or apoptosis (Kastenhuber and Lowe, 2017).

While the distinct features of the HSR, erUPR, and mitoUPR have been characterized, the existence of a discrete PQC pathway specific to the nucleus is unknown. We sought to compare the response to unfolded protein in the nucleus to that of the same unfolded protein in the cytosol. Miyazaki et al. addressed this question by monitoring a select list of genes by RT-qPCR (Miyazaki et al., 2015). In our study, we engineered mouse NIH3T3 cells to express a conditionally folded protein localized to either the nucleus or the cytosol. We then employed mRNA sequencing to capture the time-dependent global transcriptional response to compartment-specific unfolded protein. We identified a common set of stress response genes that is induced by both a nucleus-specific and cytosol-specific unfolded protein. Strikingly, we determined that only the nucleus-specific unfolded protein engages the p53 pathway and induces a transient cell cycle delay. Our work highlights the differing consequences of compartment-specific unfolded protein, particularly in the nucleus, and points to the possible role of p53 in a nucleus-specific unfolded protein response pathway in mammalian cells.

## Results

### Creating compartment-specific unfolded proteins

Destabilizing domains (DDs) are engineered single-domain proteins whose folding state depends on binding to a high affinity cell-permeable ligand (Banaszynski et al., 2006). When ligand-bound, the DD adopts a folded conformation and is metabolically stable. In the absence of the ligand, the DD partially unfolds, leading to rapid ubiquitylation and subsequent degradation by the ubiquitin-proteasome system (UPS) (Egeler et al., 2011; Iwamoto et al., 2010). The processivity of the proteasome ensures the complete degradation of the DD and any fused proteins-of-interest (Chu et al., 2013; Lee et al., 2001). In cell culture systems, ligand withdrawal is expedited by the addition of purified unliganded protein to the media (Egeler et al., 2011; Gottlieb et al., 2019; Miyazaki et al., 2016, 2012). The presence of excess extracellular protein acts a strong thermodynamic sink, facilitating intracellular ligand removal without the need for disruptive handling or media changes. In addition, this method rapidly triggers DD unfolding, leading to degradation of DDs (Chu et al., 2013; Miyazaki et al., 2015) or aggregation of the AgDD (Gottlieb et al., 2019; Miyazaki et al., 2016). Therefore, fusion of a DD to a protein-of-interest enables reversible, tunable regulation of the protein-of-interest’s expression levels in living cells through titration of ligand concentration.

Previous work from our lab used the human *FKBP1A*-derived DD fused to superfolder green fluorescent protein (DD-sfGFP) as a single species of unfolded protein to probe the cellular response (Miyazaki et al., 2015). In the present study, we began by creating DD-sfGFP constructs that differ in subcellular localization by adding localization signals to the C-terminus of sfGFP (Fig. 1A). Nuclear localization signals are short amino acid sequences that are recognized by import and export factors that actively transport proteins across the nuclear envelope (Wing et al., 2022). We named these variants by their localization: throughout the cell (non-targeted DD, ntDD), in the nucleus (NucDD), and in the cytosol (CytoDD). We then stably expressed each construct in mouse NIH3T3 fibroblasts via viral transduction. Fluorescence microscopy confirmed the expected subcellular localization of each DD-sfGFP construct in cells cultured with the stabilizing ligand Shield-1 (Fig. 1B).

**Figure 1.**
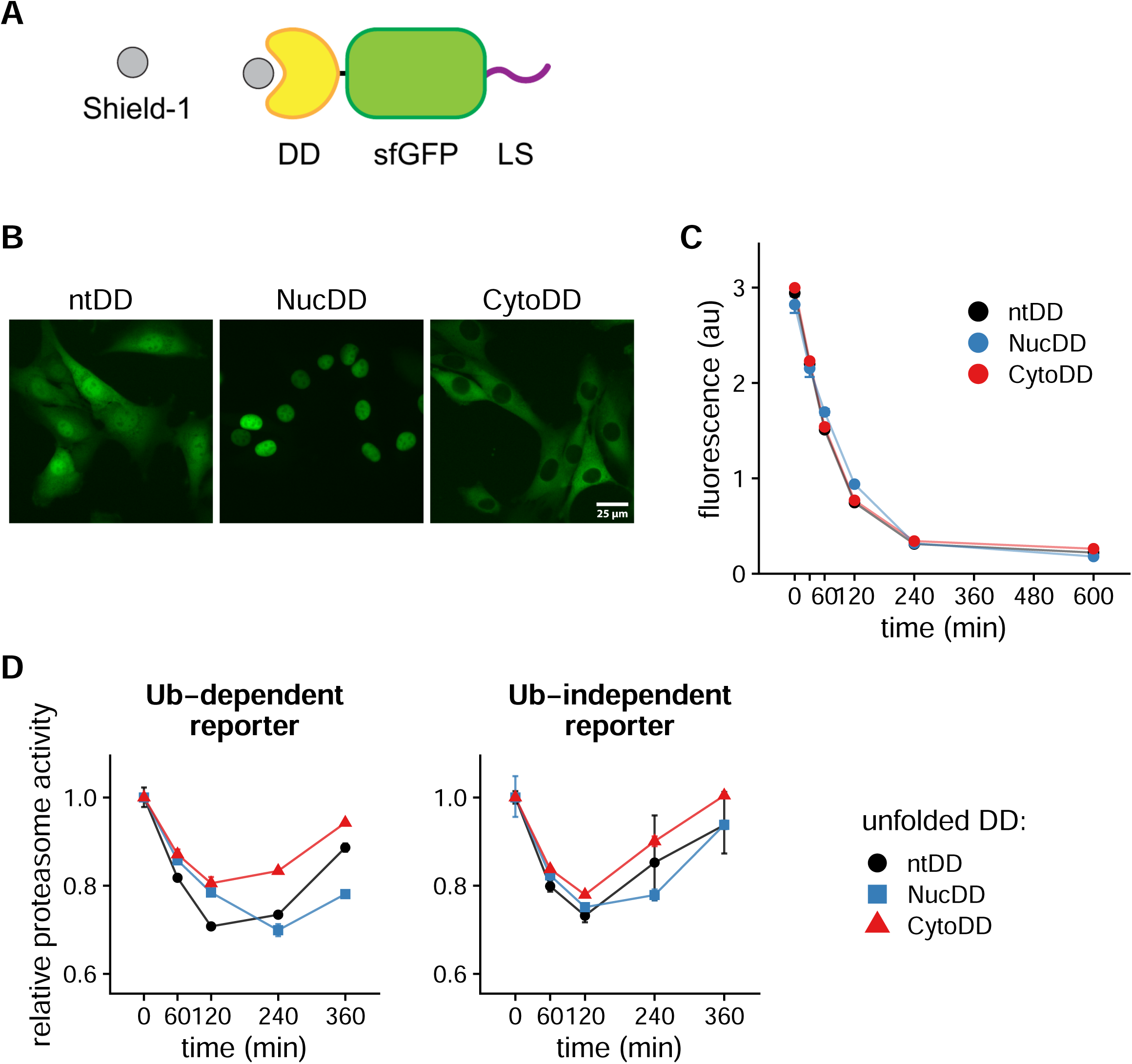
Compartment-specific unfolded DD is degraded and competes with proteasome substrates for degradation. A. Diagram of the DD-sfGFP fusion protein with C-terminal localization signal (LS). The DD is stably folded when bound to the stabilizing ligand Shield-1. The c-Myc nuclear localization signal (NLS; PAAKRVKLD) and a synthetic nuclear export signal (NES; VSSLAEKLAGLDID) were used. B. Fluorescence microscopy showing the subcellular localization of Shield-1-stabilized DD-sfGFP protein stably expressed in mouse embryonic fibroblast NIH3T3 cells. DD-sfGFP was localized by the absence of LS (left, ntDD), or by the addition of an NLS (center, NucDD) or NES (right, CytoDD) as described in A. C. DD-sfGFP fluorescence measured by flow cytometry at multiple timepoints following Shield-1 withdrawal. Each point is the mean fluorescence intensity of 20,000+ cells, and error bars indicate the standard error of the mean (sem) of 3 biological replicates. Mean fluorescence intensity is reported on a linear scale in arbitrary units (au). Dashed lines mark the half-life calculated by first order decay approximation: ntDD t_1/2_ = 57 min, NucDD t_1/2_ = 77 min, CytoDD t_1/2_ = 57 min. D. Relative proteasome activity on degron reporters calculated from the inverse (1/x) mean fluorescence intensity of mCherry-tagged degron reporters following Shield-1 withdrawal. Left, proteasome activity calculated from the effect of unfolded DD on the ubiquitin-dependent reporter (Ub^G76V^-mCh). Right, proteasome activity calculated from the ubiquitin-independent reporter (mCh-C21). Proteasome activity is reported relative to unstressed cells (0 min). Error bars indicate sem of 3 biological replicates.

We used these engineered cell lines to induce and compare the cellular responses to compartment-specific unfolded protein. First, we compared key properties of the DD constructs. In the presence of 1 µM Shield-1, the maximal mean GFP fluorescence of each construct was similar (within 6% of the mean of means; Fig. 1C, compare at 0 min). After unfolding was triggered, total GFP fluorescence levels in all 3 cell lines decreased at similar rates (half-lives ranged from 57 to 76 min) and reached near minimal levels by 240 min (Fig. 1C). These data indicate that the stability and degradation dynamics of the DD constructs in the engineered cell lines are similar despite differences in subcellular localization.

Next, we asked how the sudden appearance of unfolded DD affects UPS activity. Synthetic reporters of UPS activity have been utilized in a variety of cell culture models and model organisms (Bence et al., 2001; Bennett et al., 2005; Dantuma et al., 2000; Gierisch et al., 2020; Work and Brandman, 2021). These reporters are typically comprised of a fluorescent protein fused to a degron, which is a conserved motif that targets proteins for degradation by the proteasome (Varshavsky, 2019). Degrons are targeted to the proteasome through either ubiquitin-dependent (Ella et al., 2019) or independent (Baugh et al., 2009; Erales and Coffino, 2014) mechanisms. When UPS functionality is impaired or demand exceeds available proteolytic capacity, proteasome substrates are transiently stabilized, leading to the accumulation of the fluorescent degron reporter proteins. As UPS functionality is restored or increased, levels of these reporters return to baseline. Therefore, degron reporters provide a quantitative and dynamic readout of UPS activity in living cells.

We generated two distinct UPS activity reporters consisting of mCherry (mCh) fused to a single degron sequence. The ubiquitin-dependent degron reporter (Ub^G76V^-mCh) is rapidly poly-ubiquitylated and degraded by the proteasome through the conserved ubiquitin-fusion degradation pathway (Dantuma et al., 2000; Johnson et al., 1995). The ubiquitin-independent degron reporter (mCh-C21) utilizes a hydrophobic, proline-rich 21-amino acid sequence from the C-terminus of the transcription factor NKX3.1 that promotes proteasomal degradation without ubiquitylation (Rao et al., 2012). Although these degrons are targeted by distinct pathways, both report on proteasome activity. We first generated NIH3T3 cells stably expressing a single degron reporter by viral transduction. We then introduced each DD-sfGFP variant through a second round of viral transduction. This resulted in six cell lines co-expressing each combination of a single degron reporter and a single DD-sfGFP variant.

When cultured with Shield-1, the degron reporters are maintained at low levels by constant proteasomal degradation. We quantified ‘proteasome activity’ as the inverse (1/x) of degron reporter fluorescence levels at each time point relative to the initial unstressed state (0 min). Decreases in proteasome activity correspond to increases in the levels of the degron reporters. These events can occur due to impaired UPS function or increased demand for proteasome capacity, resulting in inefficient clearance of substrates (Hanna et al., 2007; Kandel et al., 2024). Upon exposure to any unfolded DD variant, proteasome activity on the ubiquitin-dependent degron reporter steadily decreased, reaching a minimum between 120 and 240 min (Fig. 1D, left). After 240 min, proteasome activity increased. Notably, 240 min coincides with the reduction of sfGFP-DD fluorescence signal to near basal levels (i.e., in the absence of stabilizing ligand) (Fig. 1C). We observed similar effects of the unfolded DDs on proteasome activity quantified using the ubiquitin-independent degron reporter (Fig. 1D, right). These data demonstrate that the appearance of the unfolded DD impacts UPS functionality similarly in terms of magnitude and timing despite differences in subcellular localization.

### Global transcriptional profiling reveals common and distinct features

To compare the cellular responses to compartment-specific unfolded protein, we designed an mRNA-Seq experiment using the NucDD and CytoDD cell lines. We harvested cells grown in the presence of Shield-1 (0 min), and at five additional timepoints (15, 30, 60, 120, and 240 min) after withdrawal of Shield-1 to trigger DD unfolding (Fig. 2A). We focused on these time points because the majority of unfolded DD was degraded during this interval (Fig. 1C), coinciding with a decrease in proteasome activity against the degron reporters (Fig. 1D).

**Figure 2.**
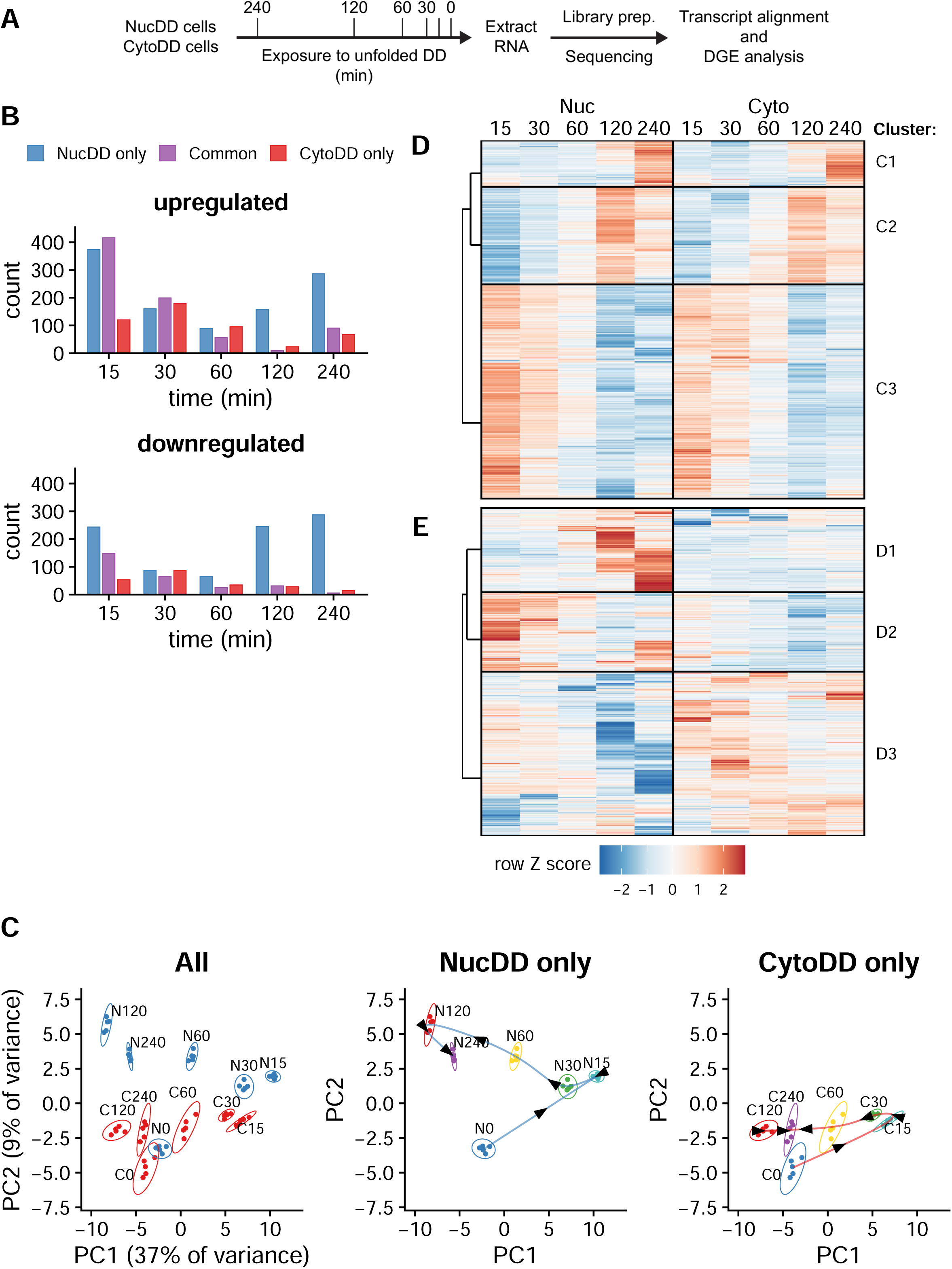
Global transcriptional profiling reveals common and distinct features. A. Schematic highlighting key experimental steps for mRNA-Seq and differential gene expression (DGE) analysis. NucDD and CytoDD cells were grown separately to log phase growth in the presence of Shield-1 in the media. Shield-1 was withdrawn to expose cells to unfolded DD for the indicated duration, then cells were harvested for total RNA. Each timepoint was prepared with 5 biological replicates. During cDNA library preparation, each sample received a unique index. Sequencing was performed on the NovaSeq 6000 platform. Transcripts were quantified using Salmon and differential gene expression analysis was performed using DESeq2. B. Quantification of differentially expressed genes (DEG) at each timepoint. Only genes that exhibited fold change (FC) >1.4 increase or decrease relative to 0 min at FDR < 0.01 were counted as significant. For each timepoint, genes were classified as ‘common’ genes (meets criteria in both NucDD and CytoDD) or ‘distinct’ genes (meets criteria in only 1 cell line). C. Principal component analysis (PCA) of all timepoints and replicates based on the top 10% most variable genes (approximately 1000 genes). Left, NucDD and CytoDD samples plotted on the same space. Center, only NucDD samples. Right, only CytoDD samples. Center and right, connecting arrows added to highlight time course progression. D. Hierarchical clustering of ‘common’ set DEGs that exhibited similar changes in response to NucDD (left) and CytoDD (right). Each row is a unique gene and gene expression is represented by a normalized row Z score relative to the 0-minute timepoint for each cell line. The dendrogram branches show the relationships of the top-level clusters. E. Hierarchical clustering of ‘distinct’ set DEGs that were differentially changed by NucDD and CytoDD, as in panel C.

We identified over 11,000 protein-coding genes with transcripts detected in both cell lines across all timepoints (Supplementary File 1). Using a fold change (FC) threshold of ± 1.5 at a false discovery rate (FDR) of < 0.01, we found 1,442 differentially expressed genes (DEGs).

Lowering the threshold to 1.2 increased the count to 5,795 DEGs. To facilitate downstream analysis, we selected an intermediate FC threshold of 1.4, resulting in 2,199 DEGs. This choice balanced the need for a sufficient number of genes to identify relevant biological patterns and pathways while focusing on the most significantly altered genes. When grouped by time point, we observed DEGs that were commonly changed in both NucDD and CytoDD cell lines as well as those that were uniquely changed in only one of the cell lines (Fig. 2B). Notably, NucDD exhibited a greater number of up- and down-regulated genes at the 120- and 240- min timepoints than CytoDD.

We applied principal component analysis (PCA) to explore the relationship between time and changes in gene expression (Fig. 2C). This analysis yielded four key observations. First, samples collected at the same time point cluster together, indicating a high concordance of data collected from each of the 5 biological replicates. Second, NucDD and CytoDD samples collected at the 0-min timepoint form neighboring clusters, indicating that the initial unstressed state is similar between these two cell lines. Third, tracing the samples in temporal sequence reveals similar cyclical paths (Fig. 2C, center and right). Fourth, although NucDD and CytoDD samples follow the same temporal sequence along PC1, they are vertically separated along PC2. This vertical separation is especially pronounced at the 120-min and 240-min timepoints. These data suggest that the cellular response to unfolded NucDD and CytoDD consists of common features that correlate with time (PC1) and distinct features that correlate with unfolded DD localization (PC2).

To identify specific patterns of expression, we employed a two-step clustering approach that simplified the complex three-dimensional relationship between DD localization, gene expression changes, and time. First, we applied k-medoids clustering (k=2) to the normalized fold change dataset. This step partitioned DEGs into two categories that we designated as either the “common set” (Fig. 2D, 1,148 genes) or the “distinct set” (Fig. 2E, 1,051 genes).

Second, we applied hierarchical clustering to each set to generate clusters, each representing a discrete pattern of expression. The common set yielded three major clusters, each exhibiting a visually distinguishable pattern shared across NucDD and CytoDD (Fig. 2D). For example, cluster C1 is enriched for genes upregulated at 240 min. The distinct set resulted in three major clusters with markedly different patterns between NucDD and CytoDD (Fig. 2E). Notably, cluster D1 contains genes that are uniquely upregulated in NucDD at the 120- and 240- min timepoints, respectively, but not in CytoDD (see Fig. 2B, top panel).

### Common response features protein quality control and oxidative stress response genes

Beginning with the common set clusters, we performed overrepresentation analysis using the Kyoto Encyclopedia of Genes and Genomes (KEGG) (Kanehisa and Goto, 2000) (Fig. 3A). Cluster C1, containing genes upregulated at 240 min in both NucDD and CytoDD, was enriched with ‘proteasome’ genes (*Psma1/5/7*, *Psmb3*, *Psmc2/6*, *Psmd1/3/11/12*, *Psme4*), ‘glutathione metabolism’ genes (*Gclm*, *Ggct*, *Gsta4*, *Gsr*, *Pgd*), and ‘protein processing in the endoplasmic reticulum’ (*Dnajb2*, *Hsph1*, *Man1a*, *Mapk9*, *Nploc4*, *Nsfl1c*, *Ube4b*, *Ubxn4*, *Vcp*, *Wfs1*). C1 contained several additional categories related to neurodegenerative disease (spinocerebellar ataxia, prion disease, amyotrophic lateral sclerosis, Huntington disease, etc.) which were predominantly comprised of UPS genes. Additionally, we performed overrepresentation analysis using Gene Ontology (GO) terms for biological processes (Ashburner et al., 2000) (Fig. 3B). Both analyses yielded similar results, particularly highlighting the common upregulation of proteasome and UPS components in cluster C1 in response to both NucDD and CytoDD.

**Figure 3.**
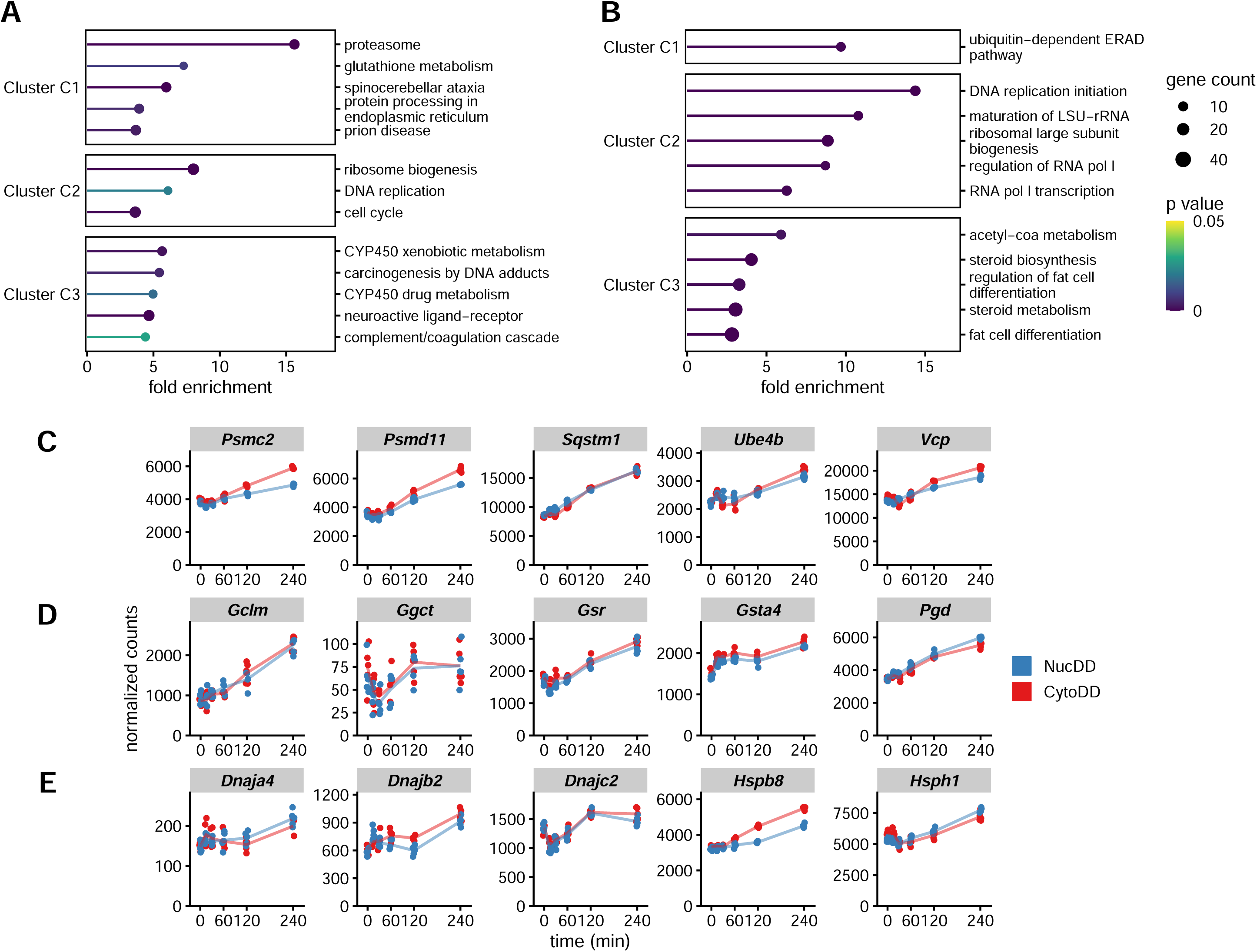
Common response features protein quality control and oxidative stress response elements. A. Enriched KEGG pathways of the common set clusters. Pathways were ranked by fold enrichment, and for visual purposes, only up to 5 significant pathways are shown (p < 0.05). Fold enrichment indicates the number of genes in each examined cluster that belong to a given pathway relative to the background. B. Enriched GO terms for biological processes of the common set cluster, as in A. C-E. Normalized counts from RNA-Seq for the indicated genes, with each point representing data from a single replicate. C. Representative ubiquitin-proteasome system genes from cluster C1. D. Representative oxidative stress response/glutathione metabolism genes from cluster C1. E. All *Hsp* and *Dnaj* chaperone proteins identified in the common set.

Using normalized row Z-scores to represent gene expression changes can obscure biologically meaningful differences in the scale of transcript abundance and magnitude of change. To confirm that our cluster analysis accurately identified comparable responses to NucDD and CytoDD, and to gain deeper insight into these DEGs, we examined the normalized counts for UPS and OSR genes in cluster C1. Normalized counts calculated by DESeq2 are well suited to comparing relative gene expression across samples (Love et al., 2014; Zhao et al., 2021). Cluster C1 UPS genes (Fig. 3C) and OSR genes (Fig. 3D) exhibited similar responses to NucDD and CytoDD when measured by normalized counts (see Fig. S3A for additional genes identified in C1 enriched pathways). These data support the robustness of our cluster and pathway enrichment analysis.

Cluster C1 overrepresentation analysis highlighted two chaperone genes, *Dnajb2* (Hsp40) and *Hsph1* (Hsp105). Chaperone proteins are critical for constitutive protein folding and refolding when faced with proteotoxic stress (Storey and Storey, 2023). As *Hsph1* has been shown to be highly inducible by heat shock in mouse models (Mahat et al., 2016), we examined the expression patterns of several chaperone proteins found in the common set of DEGs (Fig. 3E). While major inducible chaperone genes *Hspa1a/b* (Hsp70) (Radons, 2016) and *Hsp90aa1* (Hsp90) (Zuehlke et al., 2015) were found in our dataset (Supplementary File 1), they were not considered differentially expressed by our established criteria.

### Unfolded protein in the nucleus robustly induces p53 activation

We next applied overrepresentation analysis to the distinct set of DEGs (Fig. 2E). Cluster D1 genes are overrepresented by the p53 pathway (*Apaf1*, *Bbc3*, *Ccng1*, *Cdkn1a*, *Ei24*, *Fas*, *Gtse1*, *Mdm2*, *Pidd1*, *Ppm1d*, *Rrm2*, *Sesn2*, *Tnfrsf10b*, *Zfp385a*, *Zmat3*) according to both KEGG and GO analyses (Fig. 4A and B). The normalized counts for each of these genes showed that they were strongly upregulated in response to unfolded NucDD but not to CytoDD (representative genes in Fig. 4C, all genes in Fig. S4A). To validate these results, we performed RT-qPCR using *Cdkn1a* and *Mdm2* as representative p53 pathway genes (Fischer, 2017) (Fig. 4D). In a CRISPR-mediated *Trp53* (p53) knockout background NIH3T3 cell line, unfolded NucDD did not induce *Mdm2* or *Cdkn1a* (Fig. 4D) as expected (Fischer, 2017).

**Figure 4.**
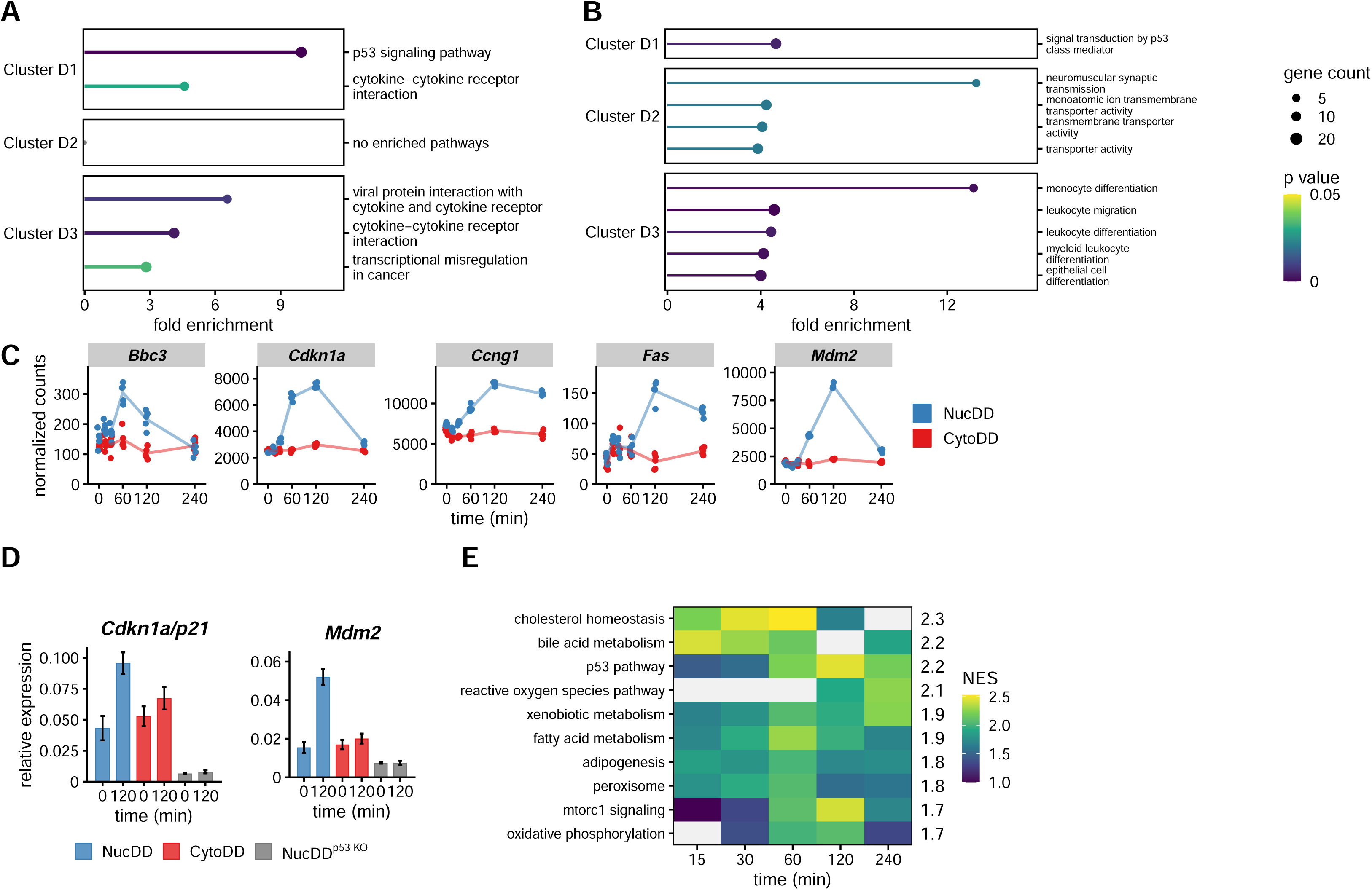
Nuclear unfolded protein robustly induces the p53 pathway. A. Enriched KEGG pathways of the distinct set clusters. See Fig. 3A for details. B. Enriched GO terms for biological processes of the distinct set cluster. C. Normalized counts for representative p53 pathway genes from cluster D1 (Fig. 2E). D. Relative gene expression of *Mdm2* and *Cdkn1a* by RT-qPCR. Expression is shown relative to *Gapdh*. Error bars indicate mean ± SEM for 3 technical replicates. E. Enriched hallmark pathways from the mouse Molecular Signature Database (MSigDb). The heatmap displays the normalized enrichment score (NES) for each timepoint. The median NES across all time points is shown on the right margin. Gray cells indicate timepoints when the indicated pathway was not significantly enriched.

This analysis of the distinct set DEGs revealed that upregulation of the p53 pathway distinguished the NucDD from the CytoDD response. However, when we examined the other enriched pathways, we did not identify any upregulated pathways that distinguished the CytoDD response from the NucDD. Therefore, we decided to concentrate our subsequent analysis on the cellular response to unfolded NucDD.

While overrepresentation analysis is useful for identifying enriched pathways within a predefined subset of genes (i.e. genes partitioned into clusters), it does not account for the strength of changes in gene expression. Gene set enrichment analysis (GSEA) is a comprehensive approach that uses a ranked list of all detected genes in the dataset to identify pathways that are enriched or depleted (Subramanian et al., 2005). To determine whether the p53 pathway is uniquely overrepresented in the cellular response to unfolded NucDD, we applied GSEA to each timepoint of the NucDD dataset using the MSigDB mouse hallmark gene set (Castanza et al., 2023; Liberzon et al., 2015; Subramanian et al., 2005). We ranked enriched pathways by highest median normalized enrichment score (NES) across all time points (Fig. 4C). Out of 26 pathways with positive median NES, the p53 pathway ranked as one of the top enriched, a result consistent with our overrepresentation analysis.

To compare the strength of pathway enrichment, we ranked the pathways by summing the normalized enrichment scores for each pathway from the 60-, 120-, and 240- min timepoints. The ‘p53 pathway’ scored highest (6.86), followed by ‘xenobiotic metabolism’ (6.25), ‘mTorc1 signaling’ (6.21), and ‘fatty acid metabolism’ (5.95). The ‘xenobiotic metabolism’ category was enriched (p < 1e-5) for glutathione metabolism/oxidative stress response genes (*Gclc*, *Gsr*, *Gss*, *Gsta1*, *Gstm4*, *Gsto1*, *Idh1*, *Pgd*). Strikingly, the ‘mTorc1 signaling’ category was enriched (p < 1e-5) for proteasome genes (*Psma3/4*, *Psmb5*, *Psmc2/4/6*, *Psmd12/13/14*, *Psme3*). Taken together, these analyses show that the p53 transcriptional program is robustly activated by the sudden appearance of unfolded protein in the nucleus, but not in the cytosol. In addition, proteasome genes and oxidative stress response genes are highly upregulated, and this is a common feature in the cellular response to both unfolded NucDD and CytoDD.

### Unfolded protein in the nucleus induces a transient p53-independent cell cycle arrest

Given the established link between p53 activation and cell cycle arrest (Kastenhuber and Lowe, 2017), we hypothesized that unfolded NucDD would induce cell cycle defects via p53 activation whereas CytoDD would not exhibit this effect. To test this, we assessed cell cycle profiles after exposure to unfolded protein by measuring total DNA content and 5-ethynyl-2′-deoxyuridine (EdU) incorporation using flow cytometry (Buck et al., 2008; Salic and Mitchison, 2008). We used an automated gating workflow to assign cell populations to specific phases of the cell cycle based on DNA- and EdU- fluorescent staining intensities (Fig. 5A).

**Figure 5.**
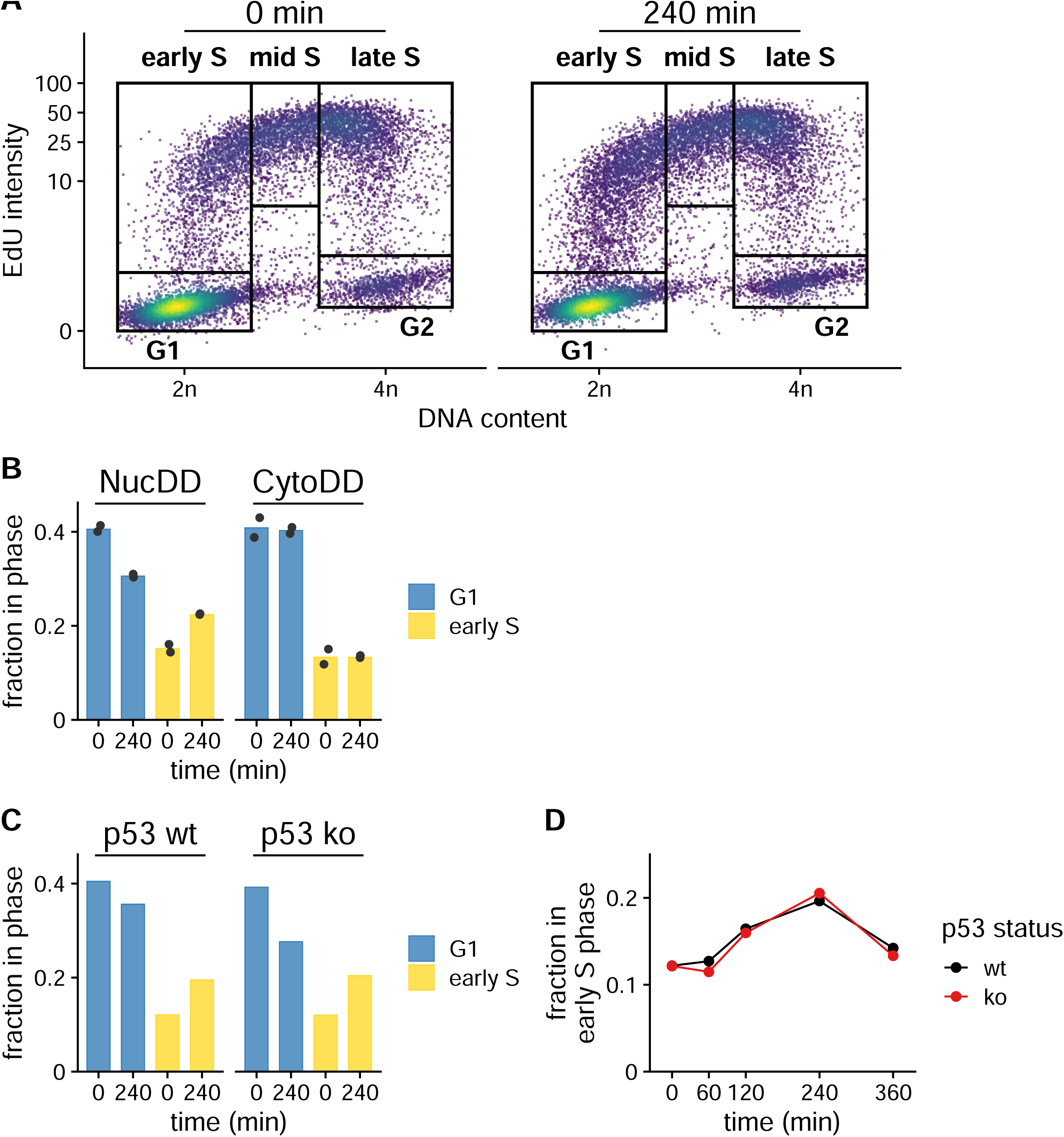
Nuclear unfolded protein induces a transient p53-independent cell cycle delay. A. Representative cell cycle analysis of total DNA content versus EdU mean intensity in NucDD cells. NucDD cells were untreated (left) or treated with ligand withdrawal for 240 min (right). At 120 min, 10 µM EdU was added to the culture medium. Boxes indicate the gating scheme used to assign cell cycle phase populations used for downstream analysis. EdU intensity is displayed as a percentage of the maximum intensity on a logarithmic scale. B. Mean fraction of NucDD and CytoDD populations gated for G1 or early S phase in untreated (0 min) or treated (240 min) cells from 2 biological samples. Data from each sample is represented as a point. C. Fraction of NucDD populations expressing either wild-type p53 or knockout gated for G1 or early S phase in untreated (0 min) or treated (240 min) cells. D. Fraction of p53^wt^ or p53^ko^ NucDD cells in early S phase treated with ligand withdrawal for the indicated times.

Strikingly, we observed that exposure to unfolded NucDD for 240 min reduced the proportion of cells in G1 and increased the population in early S phase (Fig. 5B, left). In contrast, exposure to unfolded CytoDD did not induce similar changes in G1 or early S (Fig. 5B, right). We did not observe significant changes in other phases (Fig. S5A).

Next, we tested the involvement of p53 in mediating this cell cycle arrest. We analyzed cell cycle profiles of either wild-type or CRISPR-mediated *Trp53* (p53) knockout cells expressing NucDD. Consistent with previous experiments, exposure to unfolded NucDD induced an accumulation of cells in early S phase (Fig. 5C). Unexpectedly, this result was independent of p53 status since p53 knockout cells exhibited the same accumulation in early S phase (Fig. 5C). In wild-type and knockout backgrounds, we observed that the accumulation of cells in early S phase was highest at 240 min and decreased by 360 min (Fig. 5D). Notably, the 240 min timepoint also coincides with the timing of depleted unfolded DD fluorescence signal (Fig. 1B).

### Unfolded DDs induce compartment-specific demand on proteolytic capacity

p53 is a short-lived protein and its transcriptional activity is associated with increased stability (Kastenhuber and Lowe, 2017), therefore we immunoblotted cellular lysates for levels of p53 after Shield-1 withdrawal. We observed that protein levels of p53 as well as its target gene p21 increased after 60 min of exposure to unfolded NucDD (Fig. 6A). While p53 protein levels decreased at 240 min, p21 levels remained elevated. In contrast, unfolded CytoDD did not stabilize p53 or p21 (Fig. 6A). Substituting a different fluorescent protein (mCherry) or using an orthogonal DD derived from *E. coli* DHFR (DHFRDD-sfGFP-NLS) also stabilized p53 (Fig. 6B). RNA-Seq analysis indicated that p53 mRNA was not significantly upregulated following exposure to either unfolded NucDD or CytoDD (Fig. 6C).

**Figure 6.**
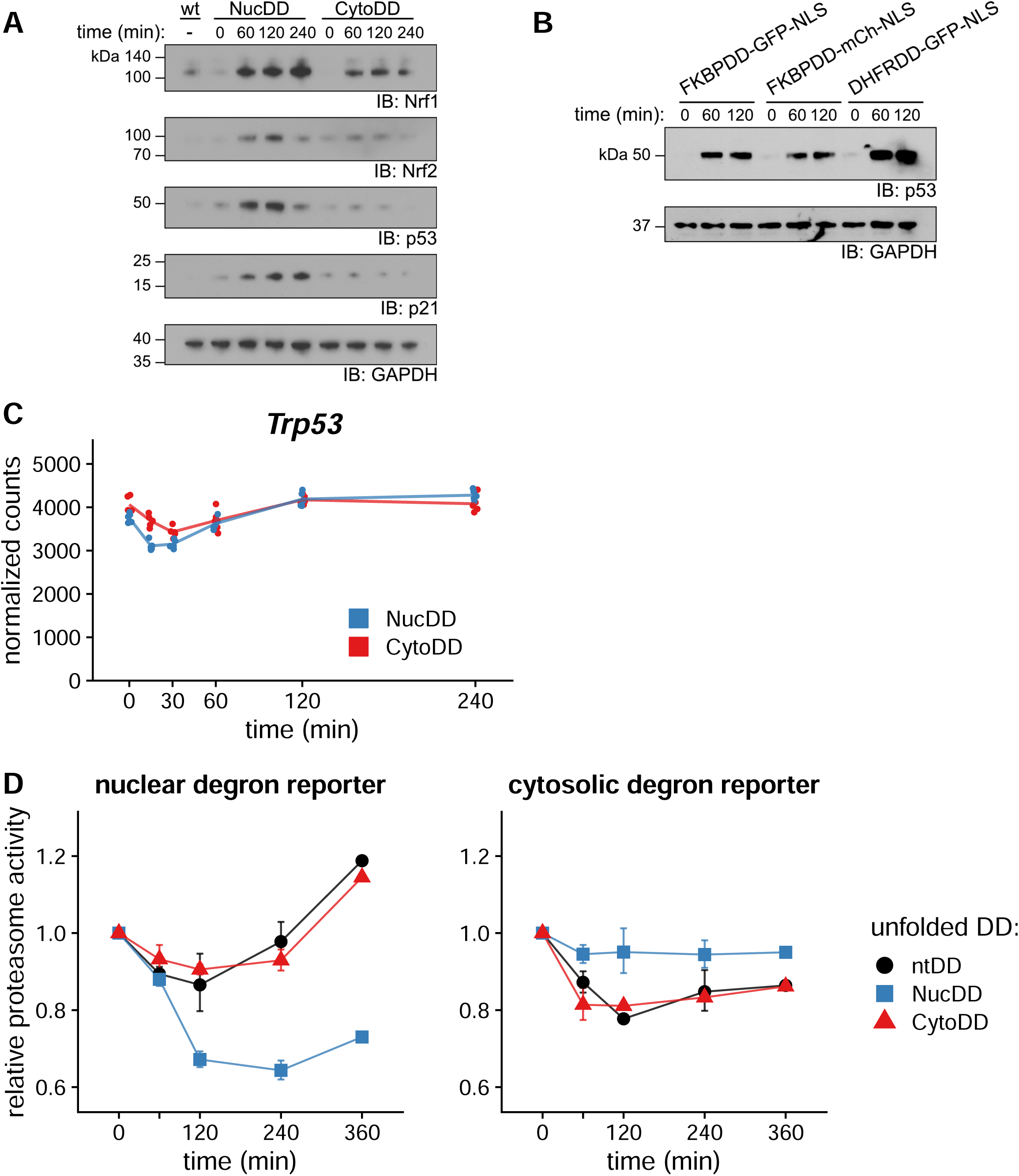
Compartment-specific increase in proteasome demand stabilizes short-lived transcription factors. A. Shield-1 was withdrawn from NucDD and CytoDD cells for the indicated times, and equal amounts of total lysate protein were immunoblotted with antibodies against Nrf1, Nrf2, p53, p21, and GAPDH. B. Nuclear DD domain variants were engineered and expressed in NIH3T3 cells. The cognate stabilizing ligand (Shield-1 for FKBPDD, TMP for DHFRDD) was withdrawn and cells were harvested at the indicated timepoints. Total lysates were immunoblotted with antibodies against p53 and GAPDH. C. Normalized counts for *Trp53* from RNA-Seq, as in Fig. 5E. D. Comparison of compartment-specific relative proteasome activity (see Fig. 1D for additional details). Ubiquitin-dependent degron reporters were localized to the nucleus or cytosol, and co-expressed with a DD-sfGFP construct. Left, proteasome activity calculated from the effect of unfolded DD on the nuclear degron reporter. Right, proteasome activity calculated from the cytosolic degron reporter. Proteasome activity is reported relative to unstressed cells (0 min). Error bars indicate sem of 3 biological replicates.

The short-lived transcription factor Nrf1 is the primary regulator of proteasome genes (Northrop et al., 2020) and Nrf2 regulates oxidative stress response genes (Ngo and Duennwald, 2022). Similar to p53, their ability to induce transcription is associated with increased stability. Since the RNA-Seq analysis showed that both NucDD and CytoDD induced gene products of these transcription factors (Ibrahim et al., 2020), we also immunoblotted for cellular levels of Nrf1 and Nrf2 following Shield-1 withdrawal (Fig. 6A). Nrf1 was stabilized by both unfolded NucDD and CytoDD. Nrf2 was stabilized by unfolded NucDD and weakly stabilized by CytoDD.

The short-lived transcription factors we identified possess degron sequences and are known to be regulated by the UPS and maintained at low levels under normal conditions (Kobayashi et al., 2004; Northrop et al., 2020; Pant and Lozano, 2014). Since the unfolded DDs are proteasome substrates and compete for proteasome capacity, we hypothesized that differences in increased stability of p53, Nrf1, and Nrf2 due to competition at the proteasome with the unfolded DD could partially be explained by localization. We wanted to test how compartment-specific proteasome capacity is affected by localized DDs. We designed localized versions of the degron reporters (Ub^G76V^-mCherry-NLS or Ub^G76V^-mCherry-NES) (Bennett et al., 2005), then co-expressed them with DDs in NIH3T3 cells. We observed that proteasome activity on the nuclear degron was more strongly affected by the presence of unfolded NucDD compared to CytoDD and ntDD (Fig. 6D, left). Conversely, proteasome activity on the cytosolic degron was more strongly affected by the presence of unfolded CytoDD and ntDD than by NucDD (Fig. 6D, right). We conclude that unfolded DD primarily attenuates proteasome capacity within the same cellular compartment.

## Discussion

The compartmentalization of eukaryotic cells, facilitated by membrane-bound organelles, confers tremendous advantages in the regulation and specialization of metabolic functions (Bar-Peled and Kory, 2022). However, this spatial separation of cellular components also introduces challenges. Each compartment requires PQC surveillance as proteins are vulnerable to misfolding due to mutations, errors during synthesis, cellular stress, and environmental conditions.

Over the last decade, researchers have made progress in characterizing PQC pathways specific to the mammalian nucleus (Azkanaz et al., 2019; Frottin et al., 2019; Guo et al., 2014; Keiten-Schmitz et al., 2020). Various experimental approaches and perturbations have facilitated discoveries and informed our current understanding of nuclear PQC mechanisms.

Acute perturbations, such as heat shock or chemical stress, rapidly induce cellular responses but are non-specific, affecting many cellular compartments and biomolecules, thus triggering a wide range of downstream effects. Alternatively, using transgenes to express mutant misfolded proteins enables greater specificity over stress localization and impacted cellular components. However, mutant transgene-induced stress is typically not acute due to the required time for transcription, translation, and accumulation of misfolded products, limiting their effectiveness for probing the immediate kinetics of stress responses. Nuclear PQC studies face an additional challenge in targeting perturbations specifically within the nucleus, while maintaining nucleocytoplasmic transport and communication. Since transcription occurs in the nucleus and protein synthesis in the cytoplasm, intact nucleocytoplasmic transport is an essential element of a coordinated cellular response. These challenges have made it difficult to investigate whether the nucleus is surveilled by a unique PQC program, analogous to the erUPR.

DDs are conditionally folded proteins with features well-suited as molecular tools to address these challenges. First, as genetically encoded transgenes, ligand-stabilized DDs can be localized to compartments like the nucleus and cytoplasm (Miyazaki et al., 2015; Sellmyer et al., 2012). Second, DD unfolding is rapidly triggered by withdrawal of the stabilizing ligand, as evidenced by their recognition and ubiquitylation by the UPS on the order of minutes (Chu et al., 2013; Egeler et al., 2011; Gottlieb et al., 2019). Third, the sudden appearance of unfolded DD elicits a cellular response in a dose-dependent manner (Miyazaki et al., 2015). This suggests that the burden created by unfolded DD is substantial enough to exceed a threshold, triggering a cellular response analogous to the erUPR. In the erUPR, this threshold is the point at which levels of misfolded proteins in the ER lumen exceed the baseline capacity of the ERAD system mitigate unfolded protein stress (Hetz and Papa, 2018). Here we leveraged these features to investigate the response to the acute appearance of unfolded protein in the nucleus of mammalian cells.

Our lab previously identified the p53 pathway as a novel component of the cellular response to unfolded DD expressed throughout the cell (Miyazaki et al., 2015). However, technical constraints limited us from determining whether unfolded DD in the nucleus and cytosol activate the p53 pathway and other identified genes to the same extent. In the present study, we designed new DD constructs that are matched in molecular size, sequential order of domains, strength of compartment-specific targeting signals, and expressed in cells at comparable total levels by fluorescence (overall lower than in Miyazaki et al. 2015, *data not shown*). These engineered improvements enabled us to differentiate between commonly induced genes and genes that are distinctly induced by NucDD and CytoDD. Importantly, we found that the p53 pathway is uniquely activated by unfolded DD in the nucleus and that it stands out as the most significantly enriched upregulated pathway.

### How does nuclear unfolded protein activate p53?

p53 is a short-lived transcription factor whose stability is tightly regulated by multiple E3 ligases (e.g. Mdm2) leading to proteasomal degradation (Kastenhuber and Lowe, 2017). The mechanisms that stabilize and activate p53 are stress-dependent. For example, phosphorylation of p53 by DNA damage response kinases stabilizes p53 by blocking recognition by E3 ligases (Abuetabh et al., 2022). We have examined the DD-expressing cells but found no evidence that the unfolded DD activates markers of the DNA damage response pathway, suggesting that DNA damage is not the driver of p53 stabilization (Miyazaki et al., 2015). In contrast, treatment of cells with proteasome inhibitors (e.g. bortezomib) stabilizes p53 regardless of post-translational modification status by blocking proteasomal degradation. The stabilization and accumulation of p53 protein promotes its nuclear occupancy and tetramerization, and tetrameric p53 binds DNA with high affinity to drive the transcription of its target genes (Chène, 2001).

We hypothesize that nuclear unfolded protein stabilizes p53 by competing for proteasomal degradation capacity in the nucleus. Our results from using localized degron reporters demonstrate that unfolded DD promotes cis-compartment degron reporter stabilization more strongly than trans-compartment degron reporters (Fig 6D). In other words, due to increased competition for available local proteolytic capacity, short-lived proteins in the same compartment as unfolded DD are not efficiently degraded and thus transiently accumulate. As a transcription factor, p53 protein is synthesized by ribosomes in the cytosol and shuttled between the nucleus and cytosol (O’Keefe et al., 2003). Notably, p53’s NES becomes hidden in the tetrameric state (Foo et al., 2007), enhancing its retention in the nucleus. Acutely interfering with nuclear proteasome capacity to degrade constitutive substrates promotes p53’s stabilization and nuclear residency, promoting tetramerization and transactivation of target genes. Conversely, when cytosolic proteasomes are overburdened by cytosolic unfolded protein, p53 is still imported into the nucleus where it can be degraded by unburdened nuclear proteasomes. Future studies are needed to test these ideas.

### Limitations and next steps

Our work reveals that p53 is sensitive to proteasome stress in the nucleus. While these studies suggest a possible role for p53 in nuclear protein quality control surveillance, it is important to acknowledge limitations stemming from the artificial nature of the DD system. Notably, it remains unclear whether the stress caused by the sudden appearance of an unfolded engineered protein is representative of natural stress. We speculate that biological stressors such as viral infection and aneuploidy might lead to a sudden accumulation of unfolded protein in the nucleus. For example, viral infection can trigger the production of viral proteins that perturb cellular proteostasis to promote aggregation (Muscolino et al., 2021). Aneuploidy can result in aberrant gene balance, resulting in the production of protein complexes with unbalanced stoichiometry of subunits and orphan proteins (Oromendia and Amon, 2014).

Mislocalized and orphan proteins, especially ribosomal proteins and proteins with intrinsically disordered domains, are known to accumulate in the nucleus and resemble misfolded proteins (Kong et al., 2021). Future work will dissect how our findings can help inform our understanding of cellular response to stress and its relevance to human health.

Our work identified a specific set of p53 genes that are induced by the unfolded DD over a period of four hours. However, not all experimentally validated p53 target genes in mouse models were upregulated according to our RNASeq analysis (Fischer, 2017). This selective gene activation highlights the complexity of p53 regulation and function. p53 regulates several transcriptional programs that can direct cell fate towards cell cycle arrest or apoptosis, survival or death (Fischer and Sammons, 2024; Kastenhuber and Lowe, 2017). There remain important open questions on how p53 integrates information from multiple signaling pathways and cellular context to direct cell fate (Fischer and Sammons, 2024; Vousden, 2000).

Our studies shed new light on the shared and distinct protein quality control landscapes of the nucleus and cytosol. In particular, we highlight the specific sensitivity of p53 to nuclear proteasome stress, suggesting that p53 may play a role in nuclear protein quality control surveillance. Future work towards understanding the mechanisms underlying these responses could provide new insights into the cellular strategies for maintaining proteostasis, especially in the nucleus where protein quality control is critical for protecting genomic integrity.

## Supplementary files

Supplementary file 1. Log2 fold change and Benjamini-Hochberg (BH) adjusted p-value data for genes identified in NucDD and CytoDD across all time points.

## Methods

### Cloning

DNA constructs were designed using SnapGene software (www.snapgene.com). Constructs were cloned into the pBMN retroviral expression vector (Nolan Lab, Stanford University) using appropriate restriction enzymes (New England Biolabs), Phusion or Q5 DNA polymerase (New England Biolabs), and NEBuilder HiFi (New England Biolabs). We used the c-Myc NLS sequence (PAAKRVKLD) for nuclear localized constructs (Ray et al. 2015). We used a synthetic NES sequence (VSSLAEKLAGLDID) for cytosolic localized constructs (Kosugi et al. 2014).

When designing DD constructs, the localization signals were fused at the C-terminus to minimize any potential effects on N-terminal DD folding. We incorporated the V206K mutation into sfGFP to promote monomerization (Zacharias et al., 2002).

### Cell culture

Mouse NIH3T3 fibroblast cells (ATCC CRL-1658) were cultured at 37°C in a standard incubator with 5% CO_2_ in Dulbecco’s Modified Eagle Medium (DMEM, Gibco) supplemented with 10% heat-inactivated adult donor bovine serum (DBS, Gibco), 100 U/mL penicillin, 100 μg/mL streptomycin, and 2 mM glutamine (PSG, Gibco). Ecotropic Platinum-E (Plat-E) retroviral packaging cells (Cell Biolabs) were cultured at 37°C in a standard incubator with 5% CO_2_ in DMEM supplemented with 10% heat-inactivated fetal bovine serum (FBS, Gibco) and PSG. Cells expressing DD constructs were cultured in the presence of the cognate stabilizing ligand: 1 µM Shield-1 for FKBPDD and 5 µM trimethoprim (TMP) for DHFRDD.

### Cell line generation

Retrovirus was produced by transfecting Platinum-E (Plat-E) cells with pBMN plasmids using TransIT-LT1 (Mirus) according to the manufacturer’s standard instructions. Viral supernatants were harvested 48 hours post-transfection and filtered (0.45 µm syringe filter, Fisher). NIH3T3 cells were incubated with retroviral supernatant supplemented with 4 µg/mL polybrene (MilliporeSigma) for 4 hours. Cells were washed once with PBS and then cultured in fresh media for 48 hours to allow for viral integration into the genome. Transduced cells were drug selected by passaging them for one week in 2 μg/ml puromycin or 5 μg/ml blasticidin.

### Destabilizing Domain (DD) unfolding

To initiate DD unfolding, we added unliganded recombinant protein purified from BL21 *E. coli* cells to the cell culture media. For FKBPDD-expressing cells, we added purified His_6_-FKBP^F36V^ in phosphate-buffered saline (PBS) to a final concentration of 5 µM. For DHFRDD-expressing cells, we added purified His_6_-ecDHFR^WT^ to a final concentration of 50 µM. The presence of excess recombinant protein in the culture media creates a thermodynamic sink that facilitates the withdrawal of Shield-1 from cells without the need for a media change (Chu et al., 2013; Miyazaki et al., 2015). Recombinant proteins were expressed and purified as previously described (Chu et al., 2013; Egeler et al., 2011; Miyazaki et al., 2015).

### Microscopy

Cells were plated on 96-well glass bottom plates (Cellvis) in DMEM lacking phenol red (Gibco) supplemented with 10% DBS and PSG. After 48 hours, live cells were imaged on a ImageXpress Micro XLS microscope (Molecular Devices) using the 10x lens (Nikon CFI Plan Fluor, NA 0.3) and Andor Zyla 4.2 sCMOS camera.

### Flow cytometry

Live cells were trypsinized, resuspended in culture media, and kept on ice before flow cytometry analysis. Fixed cells were resuspended in PBS + 1% BSA (bovine serum albumin) and kept on ice. We collected flow cytometry data at the Stanford Shared FACS Facility using the Agilent NovoCyte Quanteon and NovoCyte Penteon flow cytometers. We conducted data analysis in R using the *flowCore* package in the Bioconductor project.

### mRNA-Seq

We harvested total RNA using RNeasy kits (Qiagen), QIAShredder columns (Qiagen), and RNase-free DNase (Qiagen). Bioanalyzer (Agilent) was used to check total RNA quality and concentration. The cDNA library was prepared using the Illumina TruSeq stranded mRNA kit with 96 unique indices according to the manufacturer’s instructions. Sequencing was performed on a NovaSeq 6000 using an S2 flowcell with 2×100 bp paired-end reads.

### RNA-Seq analysis

We performed transcript mapping and quantification using Salmon (Patro et al. 2017). We conducted differential gene expression analysis using R and DESeq2 (Love et al. 2014). Genes were considered differentially expressed at absolute fold-change (FC) >1.4 and FDR controlled at a Benjamini-Hochberg (BH) adjusted p-value <0.01.

For the two-step clustering approach, we first calculated the distance metric using Dynamic Time Warping with the *dtw* R package (Giorgino, 2009), followed by k-medoids clustering. We then applied hierarchical clustering using Ward’s method.

We used the *clusterProfiler* R package (Wu et al., 2021; Yu, 2020; Yu et al., 2010) to conduct overrepresentation analysis (ORA) with KEGG and GO, and gene set enrichment analysis (GSEA) with the mouse MSigDB hallmark gene set (Castanza et al., 2023; Dolgalev, 2020; Liberzon et al., 2015; Subramanian et al., 2005). For ORA, the background gene set consisted of all detected protein coding genes in the RNA-Seq data. For GSEA, we used the Wald test statistic (log fold change divided by the standard error) calculated by DESeq2 to pre-rank input genes.

### Immunoblot

Cells were grown in 12-well plates, and washed with PBS before harvesting. Lysis buffer (1M Tris-HCl pH 8.0, 1% sodium dodecyl sulfate (SDS), 10% glycerol, protease inhibitor cocktail (MilliporeSigma)) was directly added, to the cells before scraping and heating cells to 95°C for 5 minutes. We quantified the total protein concentration of lysate using the Pierce BCA protein assay kit (ThermoFisher) and normalized all samples to 2.5-5 µg total protein. Samples were run on NuPAGE Bis-Tris precast polyacrylamide gels (Invitrogen) and transferred to PVDF membranes (MilliporeSigma) using the Trans-Blot Turbo system (Bio-Rad). We visualized bands using horseradish peroxidase(HRP)-conjugated secondary antibodies and Immobilon Western Chemiluminescent HRP Substrate (MilliporeSigma).

### Antibodies

Primary antibodies used to immunoblot against: GAPDH (1:10,000 mouse monoclonal 6C5, Santa Cruz Biotechnology), p53 (1:1000, mouse monoclonal 1C12, Cell Signaling Technology), Nrf1 (1:800, rabbit monoclonal D5B10, Cell Signaling Technology), Nrf2 (1:800, rabbit monoclonal D1Z9C, Cell Signaling Technology).

### RT-qPCR

Cells were harvested for total RNA using RNeasy kits (Qiagen), QIAShredder columns (Qiagen), and RNase-free DNase (Qiagen). For each reaction, 1.5 µg of total RNA was used for reverse transcription with random hexamer primers and the SuperScript IV First-Strand Synthesis kits (ThermoFisher) follow manufacturer’s instructions. cDNA was mixed with primers and iTaq Universal SYBR Green (Biorad) and prepared in triplicate reactions. Quantitative PCR was conducted on a LightCycler 480 Instrument II (Roche). Primers against *Gapdh* were used as the reference.

Primers used for RT-qPCR were sourced from PrimerBank (Spandidos et al., 2010): *Cdkn1a* (F: CCTGGTGATGTCCGACCTG, R: CCATGAGCGCATCGCAATC), *Mdm2* (F: TGTCTGTGTCTACCGAGGGTG, R: TCCAACGGACTTTAACAACTTCA), *Gapdh* (F: AGGTCGGTGTGAACGGATTTG, R: GGGGTCGTTGATGGCAACA)

### Cell cycle analysis

Cells were plated in 6- or 12-well plates and the culture media was refreshed the next day. When cells reached 60% confluency, Shield-1 was withdrawn at various times. 10 µM EdU (5-ethynyl-2′-deoxyuridine) was added for 1 hour before cells were trypsinized and harvested. Cells were fixed with 4% paraformaldehyde and EdU incorporation was detected using Click-iT Plus EdU Alexa Fluor 647 Flow Cytometry Assay Kit (ThermoFisher), according to manufacturer’s instructions. DNA content was detected using 1 µg/µL FxCycle Violet Stain (ThermoFisher) added to cells 30 minutes prior to analysis. Fluorescence was measured by flow cytometry.

### Data availability

RNA-Seq data generated from this study was deposited in the Gene Expression Omnibus (GEO accession no. [processing]).

## Acknowledgements

We thank Science Editors Network for reading the manuscript and providing helpful advice during manuscript preparation. We thank Drs. D. Jarosz, J. Frydman, J. Wysocka for their helpful advice and feedback. We thank Dr. T. Swigut for helpful advice with the RNA-Seq analysis. We thank Dr. N. Ratnayeke and the Meyer Lab for imaging assistance. We thank Dr. L.C. Chen for advice on experiment protocols. We thank the Stanford Shared FACS Facility (RRID: SCR_017788) for training and use of their instruments. We thank the Stanford Functional Genomics Facility for performing the library preparation and sequencing. The sequencing data was generated with instrumentation purchased with NIH grants 1S10OD02521201 and 1S10OD021763. J.H.P. was supported by the Stanford ChEM-H Chemistry/Biology Interface Predoctoral Training Program (T32 GM120007). This work was supported by the NIH (R01 GM145715 to T.J.W.).

The authors declare no competing financial interests.

## Author contributions

T.J.W. and J.H.P conceived the project and designed the experiments. J.H.P performed the experiments and data analysis. J.H.P wrote the manuscript. T.J.W. and J.H.P. edited the manuscript.

---

**Supplementary Figure 3.**
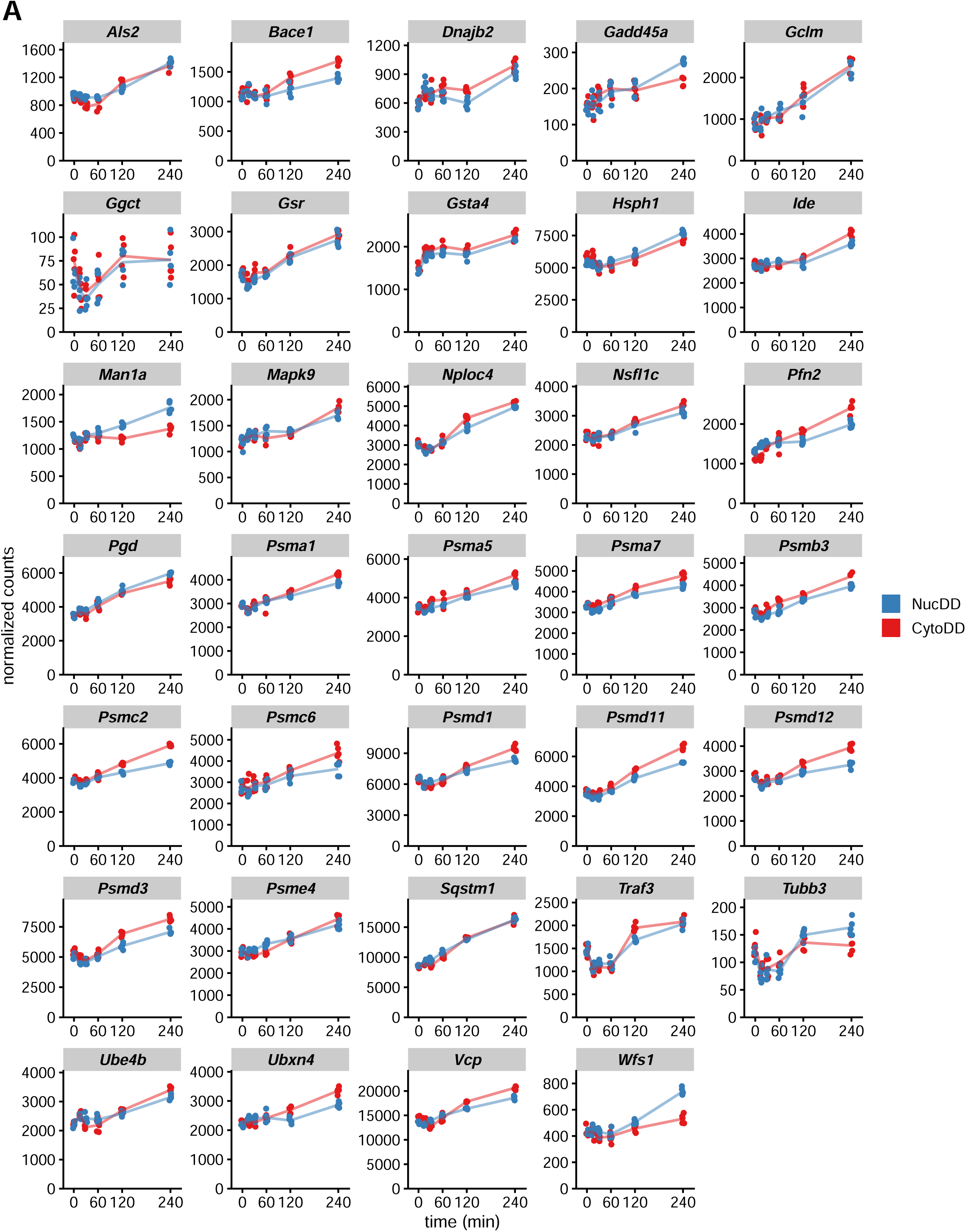
A. Normalized counts for all genes identified in significantly overrepresented KEGG pathways in cluster C1.

**Supplementary Figure 4.**
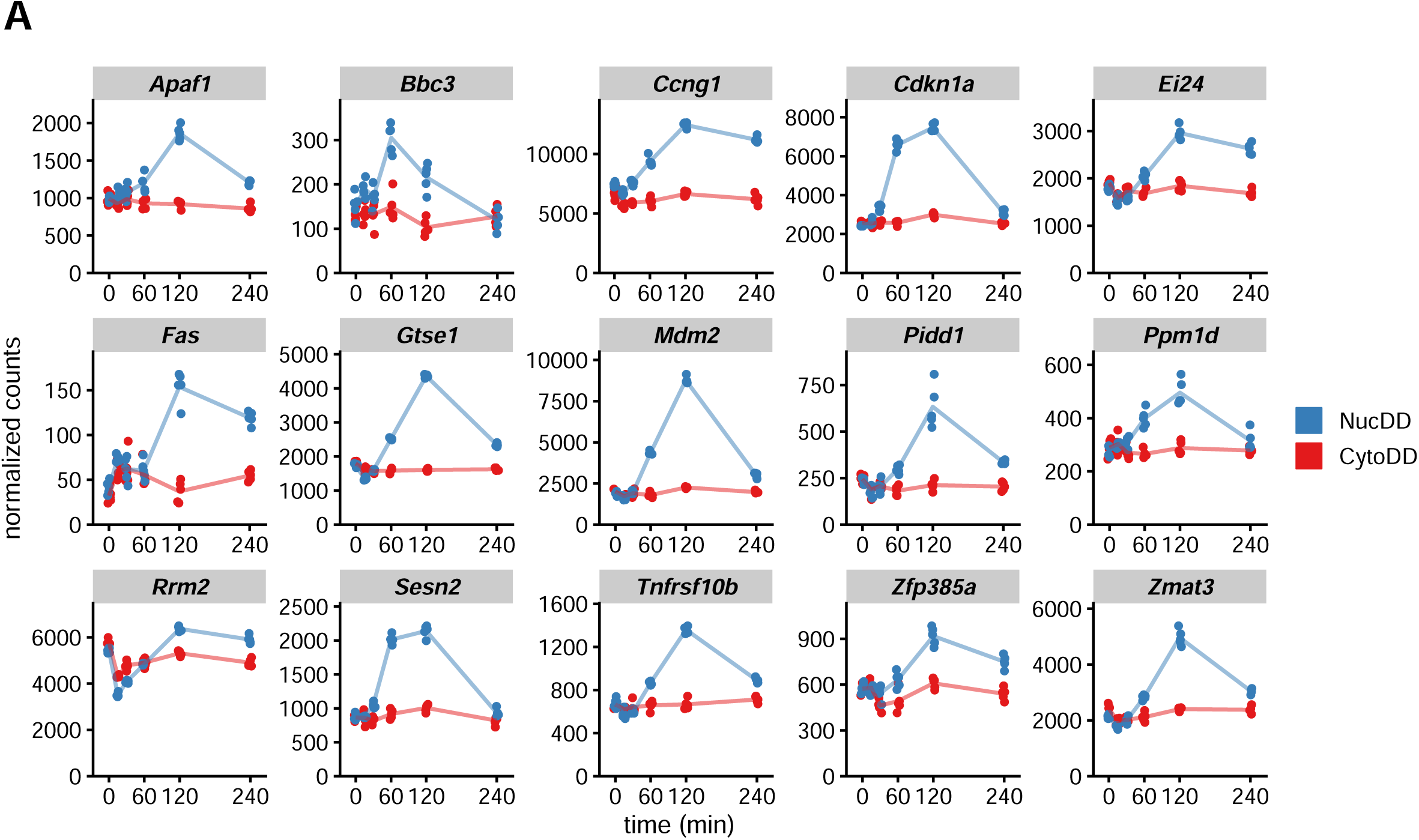
A. Normalized counts for all ‘p53 pathway’ genes identified from cluster D1 (Fig. 2E)

**Supplementary Figure 5.**
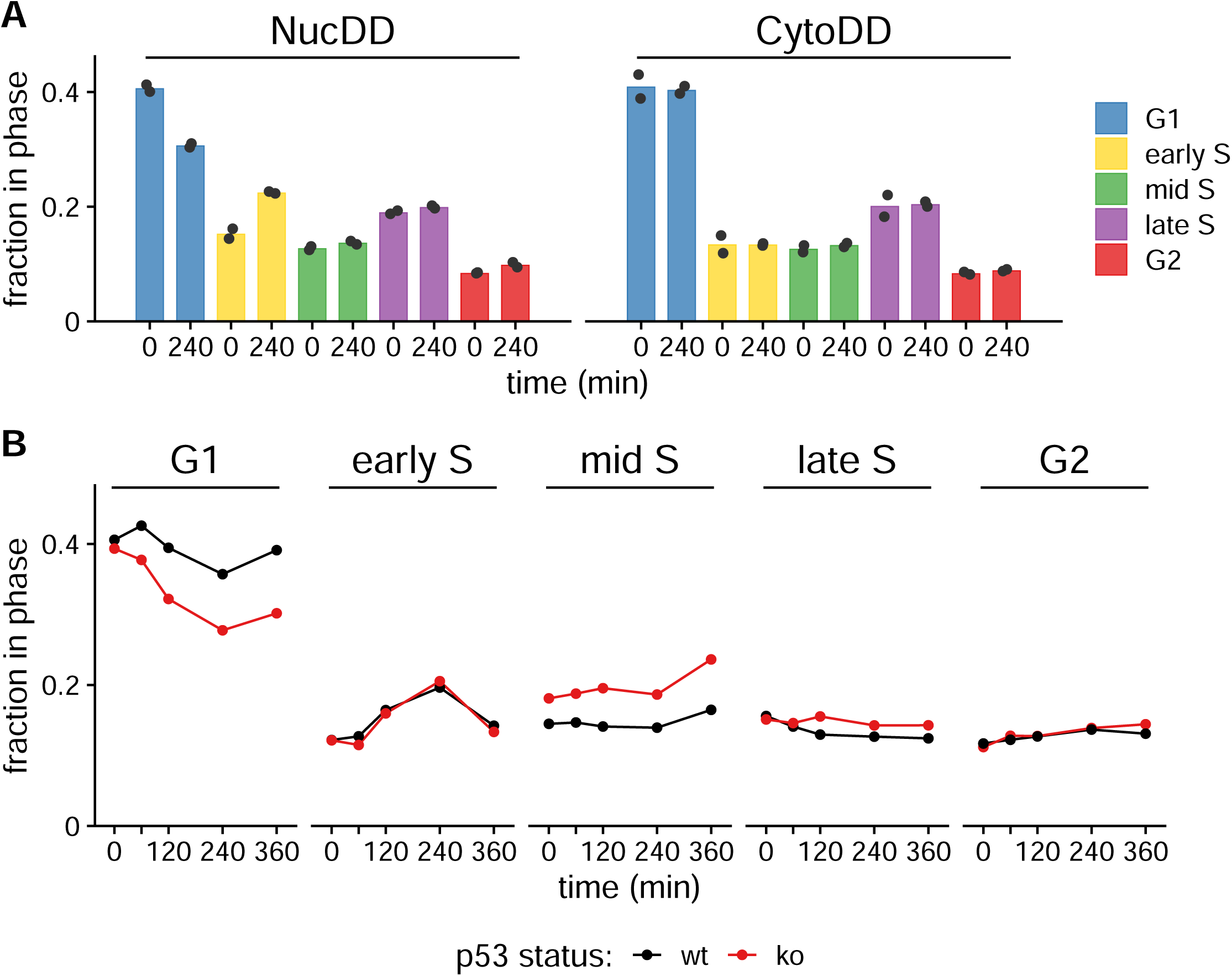
A. Mean fraction of NucDD and CytoDD populations gated for each phase in untreated (0 min) or treated (240 min) cells from 2 biological samples. Data from each sample is represented as a point. B. Fraction of p53wt or p53ko NucDD cells in each phase treated with ligand withdrawal for the indicated times.

## References

1. Abuetabh Y, Wu HH, Chai C, Al Yousef H, Persad S, Sergi CM, Leng R. 2022. DNA damage response revisited: the p53 family and its regulators provide endless cancer therapy opportunities. Exp Mol Med 54:1658–1669.

2. Ashburner M, Ball CA, Blake JA, Botstein D, Butler H, Cherry JM, Davis AP, Dolinski K, Dwight SS, Eppig JT, Harris MA, Hill DP, Issel-Tarver L, Kasarskis A, Lewis S, Matese JC, Richardson JE, Ringwald M, Rubin GM, Sherlock G. 2000. Gene ontology: tool for the unification of biology. The Gene Ontology Consortium. Nat Genet 25:25–29.

3. Azkanaz M, Rodríguez López A, de Boer B, Huiting W, Angrand P-O, Vellenga E, Kampinga HH, Bergink S, Martens JH, Schuringa JJ, van den Boom V. 2019. Protein quality control in the nucleolus safeguards recovery of epigenetic regulators after heat shock. Elife 8. doi:10.7554/eLife.45205

4. Banaszynski LA, Chen L-C, Maynard-Smith LA, Ooi AGL, Wandless TJ. 2006. A rapid, reversible, and tunable method to regulate protein function in living cells using synthetic small molecules. Cell 126:995–1004.

5. Bar-Peled L, Kory N. 2022. Principles and functions of metabolic compartmentalization. Nat Metab 4:1232–1244.

6. Baugh JM, Viktorova EG, Pilipenko EV. 2009. Proteasomes can degrade a significant proportion of cellular proteins independent of ubiquitination. J Mol Biol 386:814–827.

7. Bence NF, Sampat RM, Kopito RR. 2001. Impairment of the ubiquitin-proteasome system by protein aggregation. Science 292:1552–1555.

8. Bennett EJ, Bence NF, Jayakumar R, Kopito RR. 2005. Global impairment of the ubiquitin-proteasome system by nuclear or cytoplasmic protein aggregates precedes inclusion body formation. Mol Cell 17:351–365.

9. Buck SB, Bradford J, Gee KR, Agnew BJ, Clarke ST, Salic A. 2008. Detection of S-phase cell cycle progression using 5-ethynyl-2’-deoxyuridine incorporation with click chemistry, an alternative to using 5-bromo-2’-deoxyuridine antibodies. Biotechniques 44:927–929.

10. Buggiani J, Meinnel T, Giglione C, Frottin F. 2024. Advances in nuclear proteostasis of metazoans. Biochimie. doi:10.1016/j.biochi.2024.04.006

11. Castanza AS, Recla JM, Eby D, Thorvaldsdóttir H, Bult CJ, Mesirov JP. 2023. Extending support for mouse data in the Molecular Signatures Database (MSigDB). Nat Methods 20:1619–1620.

12. Chen B, Retzlaff M, Roos T, Frydman J. 2011. Cellular strategies of protein quality control. Cold Spring Harb Perspect Biol 3:a004374.

13. Chène P. 2001. The role of tetramerization in p53 function. Oncogene 20:2611–2617.

14. Chu BW, Kovary KM, Guillaume J, Chen L-C, Teruel MN, Wandless TJ. 2013. The E3 ubiquitin ligase UBE3C enhances proteasome processivity by ubiquitinating partially proteolyzed substrates. J Biol Chem 288:34575–34587.

15. Ciechanover A, Brundin P. 2003. The ubiquitin proteasome system in neurodegenerative diseases: sometimes the chicken, sometimes the egg. Neuron 40:427–446.

16. Cohen S, Valm AM, Lippincott-Schwartz J. 2018. Interacting organelles. Curr Opin Cell Biol 53:84–91.

17. Costa-Mattioli M, Walter P. 2020. The integrated stress response: From mechanism to disease. Science 368. doi:10.1126/science.aat5314

18. Dantuma NP, Lindsten K, Glas R, Jellne M, Masucci MG. 2000. Short-lived green fluorescent proteins for quantifying ubiquitin/proteasome-dependent proteolysis in living cells. Nat Biotechnol 18:538–543.

19. Dolgalev I. 2020. msigdbr: MSigDB gene sets for multiple organisms in a tidy data format. R package version 7.

20. Egeler EL, Urner LM, Rakhit R, Liu CW, Wandless TJ. 2011. Ligand-switchable substrates for a ubiquitin-proteasome system. J Biol Chem 286:31328–31336.

21. Ella H, Reiss Y, Ravid T. 2019. The hunt for degrons of the 26S proteasome. Biomolecules 9:230.

22. Erales J, Coffino P. 2014. Ubiquitin-independent proteasomal degradation. Biochim Biophys Acta 1843:216–221.

23. Fischer M. 2017. Census and evaluation of p53 target genes. Oncogene 36:3943–3956.

24. Fischer M, Sammons MA. 2024. Determinants of p53 DNA binding, gene regulation, and cell fate decisions. Cell Death Differ 31:836–843.

25. Foo RS-Y, Nam Y-J, Ostreicher MJ, Metzl MD, Whelan RS, Peng C-F, Ashton AW, Fu W, Mani K, Chin S-F, Provenzano E, Ellis I, Figg N, Pinder S, Bennett MR, Caldas C, Kitsis RN. 2007. Regulation of p53 tetramerization and nuclear export by ARC. Proc Natl Acad Sci U S A 104:20826–20831.

26. Frottin F, Schueder F, Tiwary S, Gupta R, Körner R, Schlichthaerle T, Cox J, Jungmann R, Hartl FU, Hipp MS. 2019. The nucleolus functions as a phase-separated protein quality control compartment. Science 365:342–347.

27. Gierisch ME, Giovannucci TA, Dantuma NP. 2020. Reporter-based screens for the ubiquitin/proteasome system. Front Chem 8:64.

28. Giorgino T. 2009. Computing and visualizing dynamic time warping alignments inR: ThedtwPackage. J Stat Softw 31. doi:10.18637/jss.v031.i07

29. Gottlieb CD, Thompson ACS, Ordureau A, Harper JW, Kopito RR. 2019. Acute unfolding of a single protein immediately stimulates recruitment of ubiquitin protein ligase E3C (UBE3C) to 26S proteasomes. J Biol Chem 294:16511–16524.

30. Guo L, Giasson BI, Glavis-Bloom A, Brewer MD, Shorter J, Gitler AD, Yang X. 2014. A cellular system that degrades misfolded proteins and protects against neurodegeneration. Mol Cell 55:15–30.

31. Hanna J, Meides A, Zhang DP, Finley D. 2007. A ubiquitin stress response induces altered proteasome composition. Cell 129:747–759.

32. Hetz C, Papa FR. 2018. The unfolded protein response and cell fate control. Mol Cell 69:169–181.

33. Hetz C, Zhang K, Kaufman RJ. 2020. Mechanisms, regulation and functions of the unfolded protein response. Nat Rev Mol Cell Biol 21:421–438.

34. Hipp MS, Kasturi P, Hartl FU. 2019. The proteostasis network and its decline in ageing. Nat Rev Mol Cell Biol 20:421–435.

35. Ibrahim L, Mesgarzadeh J, Xu I, Powers ET, Wiseman RL, Bollong MJ. 2020. Defining the functional targets of Cap’n’collar transcription factors NRF1, NRF2, and NRF3. Antioxidants (Basel) 9:1025.

36. Iwamoto M, Björklund T, Lundberg C, Kirik D, Wandless TJ. 2010. A general chemical method to regulate protein stability in the mammalian central nervous system. Chem Biol 17:981–988.

37. Johnson ES, Ma PC, Ota IM, Varshavsky A. 1995. A proteolytic pathway that recognizes ubiquitin as a degradation signal. J Biol Chem 270:17442–17456.

38. Kandel R, Jung J, Neal S. 2024. Proteotoxic stress and the ubiquitin proteasome system. Semin Cell Dev Biol 156:107–120.

39. Kanehisa M, Goto S. 2000. KEGG: kyoto encyclopedia of genes and genomes. Nucleic Acids Res 28:27–30.

40. Kastenhuber ER, Lowe SW. 2017. Putting p53 in Context. Cell 170:1062–1078.

41. Keiten-Schmitz J, Wagner K, Piller T, Kaulich M, Alberti S, Müller S. 2020. The nuclear SUMO-targeted ubiquitin quality control network regulates the dynamics of cytoplasmic stress granules. Mol Cell 79:54–67.e7.

42. Kobayashi A, Kang M-I, Okawa H, Ohtsuji M, Zenke Y, Chiba T, Igarashi K, Yamamoto M. 2004. Oxidative stress sensor Keap1 functions as an adaptor for Cul3-based E3 ligase to regulate proteasomal degradation of Nrf2. Mol Cell Biol 24:7130–7139.

43. Kong K-YE, Coelho JPL, Feige MJ, Khmelinskii A. 2021. Quality control of mislocalized and orphan proteins. Exp Cell Res 403:112617.

44. Lee C, Schwartz MP, Prakash S, Iwakura M, Matouschek A. 2001. ATP-dependent proteases degrade their substrates by processively unraveling them from the degradation signal. Mol Cell 7:627–637.

45. Liberzon A, Birger C, Thorvaldsdóttir H, Ghandi M, Mesirov JP, Tamayo P. 2015. The Molecular Signatures Database (MSigDB) hallmark gene set collection. Cell Syst 1:417–425.

46. Love MI, Huber W, Anders S. 2014. Moderated estimation of fold change and dispersion for RNA-seq data with DESeq2. Genome Biol 15:550.

47. Maciejowski J, Hatch EM. 2020. Nuclear Membrane Rupture and Its Consequences. Annu Rev Cell Dev Biol 36:85–114.

48. Mahat DB, Salamanca HH, Duarte FM, Danko CG, Lis JT. 2016. Mammalian heat shock response and mechanisms underlying its genome-wide transcriptional regulation. Mol Cell 62:63–78.

49. Miyazaki Y, Chen L-C, Chu BW, Swigut T, Wandless TJ. 2015. Distinct transcriptional responses elicited by unfolded nuclear or cytoplasmic protein in mammalian cells. Elife 4. doi:10.7554/eLife.07687

50. Miyazaki Y, Imoto H, Chen L-C, Wandless TJ. 2012. Destabilizing domains derived from the human estrogen receptor. J Am Chem Soc 134:3942–3945.

51. Miyazaki Y, Mizumoto K, Dey G, Kudo T, Perrino J, Chen L-C, Meyer T, Wandless TJ. 2016. A method to rapidly create protein aggregates in living cells. Nat Commun 7:11689.

52. Muscolino E, Luoto L-M, Brune W. 2021. Viral induced protein aggregation: A mechanism of immune evasion. Int J Mol Sci 22:9624.

53. Ngo V, Duennwald ML. 2022. Nrf2 and Oxidative Stress: A General Overview of Mechanisms and Implications in Human Disease. Antioxidants (Basel) 11. doi:10.3390/antiox11122345

54. Northrop A, Byers HA, Radhakrishnan SK. 2020. Regulation of NRF1, a master transcription factor of proteasome genes: implications for cancer and neurodegeneration. Mol Biol Cell 31:2158–2163.

55. O’Keefe K, Li H, Zhang Y. 2003. Nucleocytoplasmic shuttling of p53 is essential for MDM2-mediated cytoplasmic degradation but not ubiquitination. Mol Cell Biol 23:6396–6405.

56. Oromendia AB, Amon A. 2014. Aneuploidy: implications for protein homeostasis and disease. Dis Model Mech 7:15–20.

57. Pant V, Lozano G. 2014. Limiting the power of p53 through the ubiquitin proteasome pathway. Genes Dev 28:1739–1751.

58. Radons J. 2016. The human HSP70 family of chaperones: where do we stand? Cell Stress Chaperones 21:379–404.

59. Rao V, Guan B, Mutton LN, Bieberich CJ. 2012. Proline-mediated proteasomal degradation of the prostate-specific tumor suppressor NKX3.1. J Biol Chem 287:36331–36340.

60. Richter K, Haslbeck M, Buchner J. 2010. The heat shock response: life on the verge of death. Mol Cell 40:253–266.

61. Salic A, Mitchison TJ. 2008. A chemical method for fast and sensitive detection of DNA synthesis in vivo. Proc Natl Acad Sci U S A 105:2415–2420.

62. Santiago AM, Gonçalves DL, Morano KA. 2020. Mechanisms of sensing and response to proteotoxic stress. Exp Cell Res 395:112240.

63. Sellmyer MA, Chen L-C, Egeler EL, Rakhit R, Wandless TJ. 2012. Intracellular context affects levels of a chemically dependent destabilizing domain. PLoS One 7:e43297.

64. Shpilka T, Haynes CM. 2018. The mitochondrial UPR: mechanisms, physiological functions and implications in ageing. Nat Rev Mol Cell Biol 19:109–120.

65. Spandidos A, Wang X, Wang H, Seed B. 2010. PrimerBank: a resource of human and mouse PCR primer pairs for gene expression detection and quantification. Nucleic Acids Res 38:D792–9.

66. Storey JM, Storey KB. 2023. Chaperone proteins: universal roles in surviving environmental stress. Cell Stress Chaperones 28:455–466.

67. Subramanian A, Tamayo P, Mootha VK, Mukherjee S, Ebert BL, Gillette MA, Paulovich A, Pomeroy SL, Golub TR, Lander ES, Mesirov JP. 2005. Gene set enrichment analysis: a knowledge-based approach for interpreting genome-wide expression profiles. Proc Natl Acad Sci U S A 102:15545–15550.

68. Varshavsky A. 2019. N-degron and C-degron pathways of protein degradation. Proc Natl Acad Sci U S A 116:358–366.

69. Vousden KH. 2000. P53. Cell 103:691–694.

70. Wing CE, Fung HYJ, Chook YM. 2022. Karyopherin-mediated nucleocytoplasmic transport. Nat Rev Mol Cell Biol 23:307–328.

71. Wolff S, Weissman JS, Dillin A. 2014. Differential scales of protein quality control. Cell 157:52–64.

72. Work JJ, Brandman O. 2021. Adaptability of the ubiquitin-proteasome system to proteolytic and folding stressors. J Cell Biol 220. doi:10.1083/jcb.201912041

73. Wu T, Hu E, Xu S, Chen M, Guo P, Dai Z, Feng T, Zhou L, Tang W, Zhan L, Fu X, Liu S, Bo X, Yu G. 2021. clusterProfiler 4.0: A universal enrichment tool for interpreting omics data. Innovation (Camb) 2:100141.

74. Yang Y, Guo L, Chen L, Gong B, Jia D, Sun Q. 2023. Nuclear transport proteins: structure, function, and disease relevance. Signal Transduct Target Ther 8:425.

75. Yu G. 2020. Gene Ontology Semantic Similarity Analysis Using GOSemSim. Methods Mol Biol 2117:207–215.

76. Yu G, Li F, Qin Y, Bo X, Wu Y, Wang S. 2010. GOSemSim: an R package for measuring semantic similarity among GO terms and gene products. Bioinformatics 26:976–978.

77. Zacharias DA, Violin JD, Newton AC, Tsien RY. 2002. Partitioning of lipid-modified monomeric GFPs into membrane microdomains of live cells. Science 296:913–916.

78. Zhao Y, Li M-C, Konaté MM, Chen L, Das B, Karlovich C, Williams PM, Evrard YA, Doroshow JH, McShane LM. 2021. TPM, FPKM, or Normalized Counts? A Comparative Study of Quantification Measures for the Analysis of RNA-seq Data from the NCI Patient-Derived Models Repository. J Transl Med 19:269.

79. Zuehlke AD, Beebe K, Neckers L, Prince T. 2015. Regulation and function of the human HSP90AA1 gene. Gene 570:8–16.

